# Capillary and EndoMT-derived endothelial cells cooperatively form the angiogenic core niche in infantile hemangioma

**DOI:** 10.64898/2026.07.30.741725

**Authors:** Xiaoxia Gong, Zhaoshui Li, Lei Yang, Zitong Chen, Dianjin Xing, Lamei Yang, Luyao Wang, Junhui He, Pengjing Yu, Yaxin Niu, Liang Wang, Haiyan Zhang, Kaihui Zhang, Dan Song, Zhaojian Liu, Lei Guo

## Abstract

Infantile hemangioma (IH), the most common benign vascular tumor in infants, is characterized by the aberrant proliferation of hemangioma endothelial cells (HemECs) and excessive angiogenesis. However, it is unclear how the tumor microenvironment sustains these active angiogenic signals. We performed single-cell RNA sequencing and spatial transcriptomic to profile 30 samples from proliferating IH, involuting IH, and normal skin. The results indicated that HemECs comprised distinct subpopulations. Among them, capillary endothelial cells (Cap.ECs) had an immature phenotype and functioned as the core drivers of IH. The endothelial-to-mesenchymal transition endothelial cells (endoMT-ECs) secreted collagen and engaged in CD44-mediated crosstalk with Cap.ECs to construct the angiogenesis-active niche in the tumor microenvironment (TME) of proliferating IH. Blockade of CD44-mediated intercellular communication effectively suppresses the angiogenesis of HemECs and promotes tumor regression. Beyond the endothelial layer, pericytes tightly enveloped Cap.ECs, with *SERPINE2^+^* subtypes sustaining their angiogenic activity via ANGPT signaling. Pseudotime trajectory analysis revealed that the mesenchymal stem cells (MSCs)-enriched niche serves as a progenitor pool for Cap.ECs in proliferating IH. Our findings thus revealed that IH progression relies on CD44-mediated endoMT-EC–Cap.EC crosstalk and on both pericyte and MSC niches to sustain angiogenesis.

**HIGHLIGHTS:**

1. Capillary endothelial cells drive infantile hemangioma progression, with pericytes tightly ensheathing them.
2. CD44-mediated capillary and endoMT-derived endothelial cells crosstalk construct the angiogenic-active niche in the tumor microenvironment of proliferating infantile hemangioma.
3. The MSC-enriched niche provides a progenitor pool for the capillary endothelial cells in proliferating infantile hemangioma.

## INTRODUCTION

Infantile hemangioma (IH), the most common benign vascular tumor in infants, is characterized by the rapid proliferation of tissue followed by a spontaneous involution that is often incomplete [1]. Although many lesions self-resolve, some require interventions to prevent severe complications, such as disfigurement, ulceration, or airway obstruction [2, 3]. The current first-line treatments, primarily non-selective beta-blockers, can have adverse cardiovascular effects and require long-term administration. It is therefore necessary to have a better understanding of the pathogenesis of IH to develop novel targeted and safe therapies

Earlier studies identified a relationship of the clonal expansion of hemangioma endothelial cells (HemECs) with hypoxia-driven stem cell differentiation during the pathogenesis of IH [4]. Although glucose transporter type 1 is an established marker for HemECs [5], the increased level of this marker alone cannot explain pathogenesis or tumor dynamics. Angiogenesis drives the early proliferation phase of tumor growth [6], but the mechanism by which the tumor microenvironment (TME) sustains excessive vascular growth is largely unknown. For example, there is an incomplete understanding of the interactions of the heterogeneous subtypes of endothelial cells and surrounding stromal cells during the pathogenesis of IH.

In the present study, we used single-cell RNA sequencing (scRNA-seq) [7] and spatial transcriptomic [8] to profile samples from proliferating IH, involuting IH, and normal skin. We focused on the functional states and hypoxia-induced onset of the differentiation of HemECs. Additionally, we mapped the spatial interactions of cells within the angiogenic niche, and examined the effect of key signaling pathways on the interactions of specific subpopulations of endothelial cells with one another and with pericytes, which have a high abundance in IH. The ultimate aim is to characterize the coordinated multi-cellular network that sustains tissue proliferation during IH and to identify pathogenic pathways that can potentially be targeted by novel therapeutic targets.

## RESULTS

### HemECs have a dual malignant-cell like and immunosuppressive phenotype during the pathogenesis of IH

We initially characterized the intracellular and extracellular landscape of IH by use of scRNA-seq on 15 tissue samples, 6 from proliferative IH (Pro.IH), 4 from involutional IH (Inv.IH), and 5 from normal skin (Nor.Skin). This led to the identification of 172,683 cells in nine major lineages (Fig. 1A-C; Supplementary Fig. 1A-B; Supplementary Table 1). The results showed that disease progression was associated with a profound alteration of cellular composition. In contrast to normal skin, IH lesions had a depletion of epithelial lineages and an abundance of pericytes, fibroblasts, macrophages, and HemECs. The abundance of HemECs peaked during the proliferation phase and decreased sharply during the involution phase (Supplementary Fig. 1C). We applied CosMx® SMI to 399 fields of view in another 15 samples to characterize the tissue architecture (Fig. 1A). The resulting spatially resolved atlas of 634,558 cells confirmed the population dynamics identified by scRNA-seq and also showed that a high density of pericytes surrounded HemECs (Supplementary Fig. 1D-I). This is consistent with the clinical characteristics of IH, in which HemECs drive rapid vascular growth and pericytes tightly envelope HemECs to establish an angiogenic niche.

**Figure 1.**
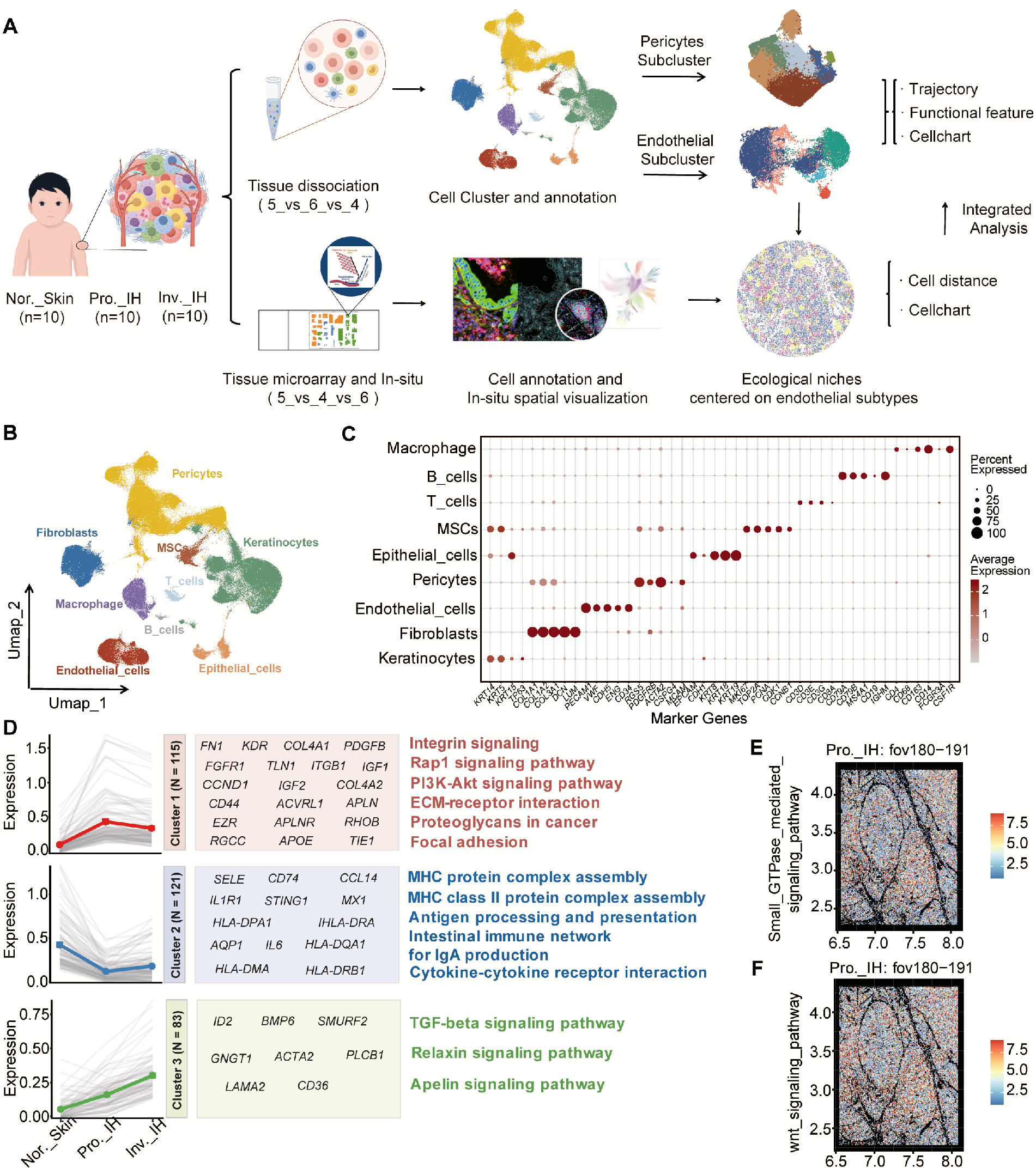
HemECs have a dual pro-angiogenic and immunosuppressive. phenotype during the pathogenesis of IH. (A) Study design. (B) Uniform manifold approximation and projection (UMAP) plot, showing 172,683 single cells (9 major cell types) from Nor.Skin, Pro.IH, and Inv.IH samples. (C) Expression of canonical markers for each major cell type. (D) Analysis of gene expression dynamics and associated GO enrichment among HemECs across sequential stages from Normal Skin (Nor. Skin) through proliferative infantile hemangioma (Pro. IH) to Involuting Infantile Hemangioma (Inv. IH). (E, F) Spatial in situ representation of activity scores for small GTPase mediated signaling pathway (E) and wnt signaling pathway (F) in the proliferative infantile hemangioma (Pro. IH) tissue.

Given the central role of HemECs and their spatial distribution, we examined the transcriptional dynamics of these cells during the different phases of IH. Clustering analysis demonstrated that three distinct gene modules were associated with the changes in tissue phenotype (Fig. 1D; Supplementary Table 2). Gene set 1 was transiently upregulated only during the proliferation phase and was highly enriched for tumor-like and pro-angiogenic pathways (e.g., PI3K-Akt, focal adhesion) (Fig. 1D; Supplementary Fig. 2A-B), consistent with the spatial pattern of *in situ* expression (Fig. 1E-F). Gene set 2, which regulates innate immunity and antigen presentation, was actively suppressed during proliferation, but was restored during involution (Fig. 1D; Supplementary Fig. 2A-B). Gene Set 3 maintained a persistently elevated expression profile across the proliferative and involuting phases of IH, with significant enrichment in the Apelin and TGF-β signaling pathways (Fig. 1D; Supplementary Fig. 2A-B). Together, these results demonstrate that proliferating IH is driven by the dual-phenotypic state of HemECs, which have increased angiogenesis and decreased immune regulation.

### Cap.ECs act as multipotent progenitors and the central hub of functionally distinct niches that orchestrate the IH microenvironment architecture

To determine the nature of the functional heterogeneity of HemECs that drives the progression of IH, we classified these cells into five distinct lineages: capillary (Cap.ECs), arterial (Art.ECs), venous (Ven.ECs), lymphatic (Lym.ECs), and pendothelial-to-mesenchymal transition endothelial cells (endoMT-ECs) (Fig. 2A-C; Supplementary Fig. 3A; Supplementary Table 4). Cap.ECs and endoMT-ECs were more abundant in proliferating IH tissues, consistent with the results from multiplex immunofluorescence (mIF) (Fig. 2B; D-F). Functional enrichment analysis revealed clear differences in activity, in that Cap.ECs had activation of core pro-angiogenic pathways (HIF-1, VEGF), whereas endoMT-ECs functioned in structural support through integrin and extracellular matrix (ECM) signaling (Fig. 2H; Supplementary Fig. 3B). Cap.ECs also had an immature, naive phenotype with pronounced tumor stemness (*ROR1^+^*, *NOX4^+^*) and increased expression of genes that function in embryogenesis (*DLK1*, *GAP43*) (Fig. 2G, Supplementary Fig. 3C-E). The endoMT-ECs were a transition state of Cap.ECs, in that they co-expressed canonical capillary markers (e.g., *RGCC*, *PECAM1/CD31*) and pericyte markers (e.g., *RGS5*, *PDGFRB*) (Fig. 2C).

**Figure 2.**
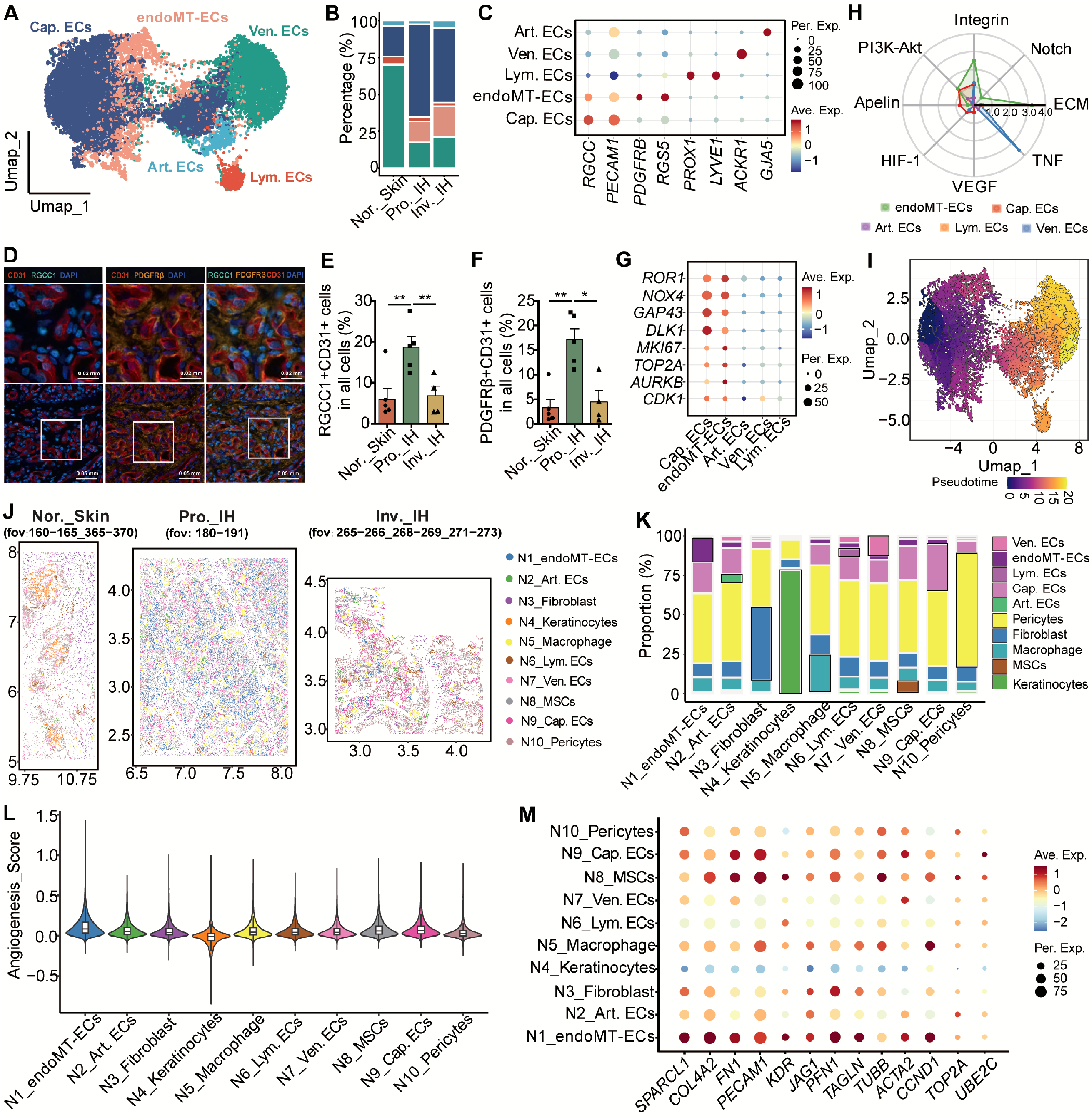
Cap.ECs act as multipotent progenitors and the central hub of functionally distinct niches that orchestrate the IH microenvironment architecture. (A) UMAP plot showing the re-clustering of 14,464 endothelial cells from Nor.Skin, Pro.IH, and Inv.IH samples. (B) The proportions of different subsets of endothelial cell subsets in Nor.Skin, Pro.IH, and Inv.IH. (C) Expression of canonical markers in each subset of endothelial cells. (D-F) mIF staining validation of Cap. ECs and endoMT-ECs in Nor. Skin, Pro. IH, and Inv. IH tissues. (D) Representative staining images, with DAPI-stained nuclei (blue). Scale bar: 50 μm. (E) Co-expression of RGCC and CD31. (F) Co-expression of PDGFRB and CD31. (G) Dot plots showing the expression levels of cancer stemness and cell proliferation marker genes for each EC subset. (H) Pathway enrichment scores for each subset of endothelial cells. (I) Single-cell trajectory analysis of endothelial cell subsets. (J) Spatial mapping of the ten tissue niches (N1–N10) identified by k-means clustering in representative tissue sections from Nor.Skin, Pro.IH, and Inv.IH. (K) Proportions of different types of cells in the ten niches. (L) Violin plot showing the differences in angiogenesis score across each niche. (M) Expression of pro-angiogenic genes in Cap.ECs from the ten niches.

Analysis of single-cell trajectories revealed a trans-differentiation from immature Cap.ECs to other endothelial states, including endoMT-ECs (Fig. 2I; Supplementary Fig. 4A-D). During this pseudotime, there was a functional shift of this trajectory in which early-stage cells had strong pro-angiogenic program (HIF-1/VEGF), and this changed to vascular-stabilizing (ECM) and immunomodulatory (antigen presentation) programs at the later stages (Supplementary Fig. 4E-F). These results indicate that Cap.ECs act as multipotent progenitors that drive the initial vascularization, and that functional switching is responsible for structural remodeling during the later phases of angiogenesis.

To understand the functions of these subtypes of HemECs within intact tissue, we integrated our scRNA-seq signatures with spatial transcriptomics data. The clustering of spatial regions according to cellular co-localization enabled classification of ten spatial niches in the IH microenvironment (Fig. 2J-K; Supplementary Fig. 5A). Thus, IH lesions had enrichment of four pathological niches: a endoMT-ECs-enriched niche (N1), a mesenchymal stem cell (MSC)-enriched niche (N8), a Cap.EC-enriched niche (N9), and a macrophage-enriched niche (N5) (Fig. 2J, Supplementary Fig. 5B-C). Notably, all four of these niches had a dense infiltration of pericytes that surrounded the Cap.ECs (Fig. 2J-K, Supplementary Fig. 5D). Pathway analysis of these niches showed that N9 (Cap.ECs) drove proliferation, with the highest activity in cell migration and proliferation; N8 (MSCs) had robust angiogenic signaling; and N1 (endoMT-ECs) had significant deposition of ECM and the highest angiogenesis score (Fig. 2L, Supplementary Fig. 6A). Moreover, the Cap.ECs in these niches, particularly in N1 (endoMT-ECs), had upregulation of genes that functioned in angiogenesis (*PECAM1*, *KDR*), cytoskeleton formation (*ACTA2*), and cell-cycle (*CCND1*) (Fig. 2M; Supplementary Fig. 6B). This tumor-like, pro-angiogenic profile indicates that Cap.ECs are the key driver to form the angiogenic microenvironment of IH.

### COLLAGEN-CD44 mediates EndoMT-ECs and Cap.ECs crosstalk to form an angiogenic niche in the TME

Because the EndoMT-ECs in N1 regulate deposition of ECM and angiogenesis (Fig. 2L-M), we investigated the spatial characteristics of these intercellular interactions. An *in situ* analysis and quantitative distance measurements demonstrated that EndoMT-ECs tightly surrounded Cap.ECs, and that this was the closest intercellular interaction within the microenvironment (Fig. 3A, B). Cell-cell communication analysis identified the COLLAGEN pathway as the dominant hub (Fig. 3C, D; Supplementary Fig. 7A). During the proliferation phase, EndoMT-ECs and Cap.ECs had increased interactions mediated by COL4A1–CD44 signaling, and this interaction decreased markedly during the involution phase (Fig. 3E; Supplementary Fig. 7B, C). Multiplex immunofluorescence microscopy of clinical samples confirmed that COL4A1 and its receptor (CD44) had high expression in this perivascular niche, suggesting that matrix-driven paracrine signaling promotes angiogenesis (Fig. 3F-H).

**Figure 3.**
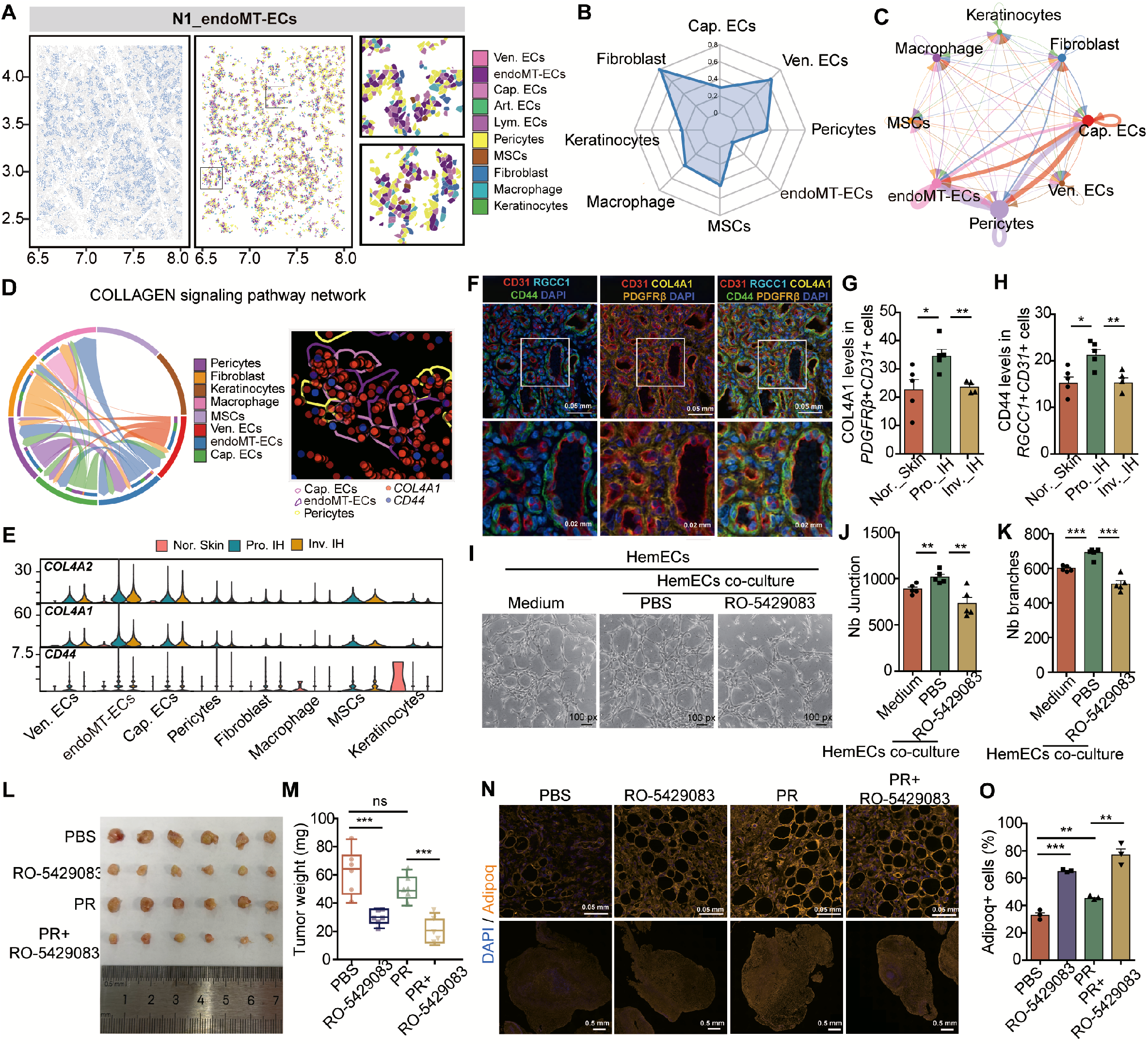
COLLAGEN-CD44 mediates endoMT-ECs and Cap.ECs crosstalk to form an angiogenic niche in the TME. (A) Spatial distribution of N1 (Peri.EC) in serial tissue sections from Pro.IH. (B) Mean quantitative distance (axis length) of different cells to Cap.ECs. (C) Cell-cell communication networks in N1 (Peri.ECs). (D) Chord diagram, showing the strength of communication of the COLLAGEN signaling pathway in N1 (Peri.ECs) (left), and in situ expression of key communication molecules (right). (E) Expression of collagen ligands and the CD44 receptor in different types of cells. (F) Representative mIF images, showing co-staining of proteins in Pro.IH tissue. (G-H) Quantification of COL4A1 and CD44 in Nor.Skin, Pro.IH, and Inv.IH. (I) Representative images of tube formation under different conditions. Scale bar: 100 μm. (J-K) Network branching and quantification of Junctions and Branches under different conditions. (L-M) Representative images of tumors (L) and tumor weights (M) in nude mice with hemangioma xenografts that received vehicle (control), CD44-blocking antibody, low-dose propranolol, or combination therapy. (N) Representative images of CD31-stained tumor sections showing vascular density in the four groups. (O) Quantification of microvessel density in the four groups.

To determine whether this spatial crosstalk promotes the formation of vessels, we performed co-culture assays with HemECs. The results indicated that HemECs increased the angiogenesis of neighboring endothelial cells *via* paracrine signaling, and that blockage of CD44 abolished this effect (Fig. 3I-K). In addition, *in vivo* analysis showed that selective inhibition of CD44 decreased the growth and vascular density of the hamangioma, but did not affect wound healing (Fig. 3L-O). Application of this targeted blockade with low-dose propranolol (the standard treatment for IH) amplified this effect. In particular, when initiated during the early proliferation phase, this treatment induced regression that remained after treatment cessation, indicating durable reprogramming of the TME (Fig. 3L-O).

These findings indicated that EndoMT-ECs provide physical and molecular support to Cap.ECs *via* a specific COLLAGEN-CD44 paracrine loop, and that blockage of this signaling disrupted the pathological vascular network. These findings have potential implications for the development of a novel therapeutic strategies for IH.

### HIF2A activates the transcription of CD44 in Cap.ECs

Because CD44 is a critical sensor of the TME in Cap.ECs, we examined the regulation of its transcription. We therefore measured the dynamics of transcription factors (TFs) in Nor.Skin, Pro.IH and Inv.IH (Supplementary Table 5). The results showed that HIF2A was the most significantly upregulated TF in proliferative Cap.ECs, with expression levels roughly twofold higher than in controls (Fig. 4A). Our analysis predicted that this TF regulated over 200 genes, including a marker of IH (*SLC2A1*) and a pro-angiogenic effector (*PECAM1/CD31*) (Supplementary Table 6). Analysis of the CD44 signaling network showed that ten TFs, including HIF2A, had high expression in proliferative Cap.ECs (Fig. 4B). Moreover, the upregulation of HIF2A had a strong positive correlation with the levels of CD44 and genes downstream of CD44 signaling (Fig. 4C, Supplementary Fig. 8A), and was highly specific to Cap.ECs (Fig. 4D, E). This relationship was not present in other subtypes of endothelial cells (Supplementary Fig. 8B-E). We confirmed that this regulation was direct based on public ChIP-seq datasets, which identified a HIF2A binding peak in the *CD44* promoter (Fig. 4F). This finding was also supported by structural modeling, which indicated high binding affinity of the HIF2A protein and the *CD44* DNA motif (Fig. 4G). In agreement, the *in silico* virtual knockout of *HIF2A* severely disrupted organization of the ECM and CD44 signaling (Fig. 4H, Supplementary Fig. 8F, G).

**Figure 4.**
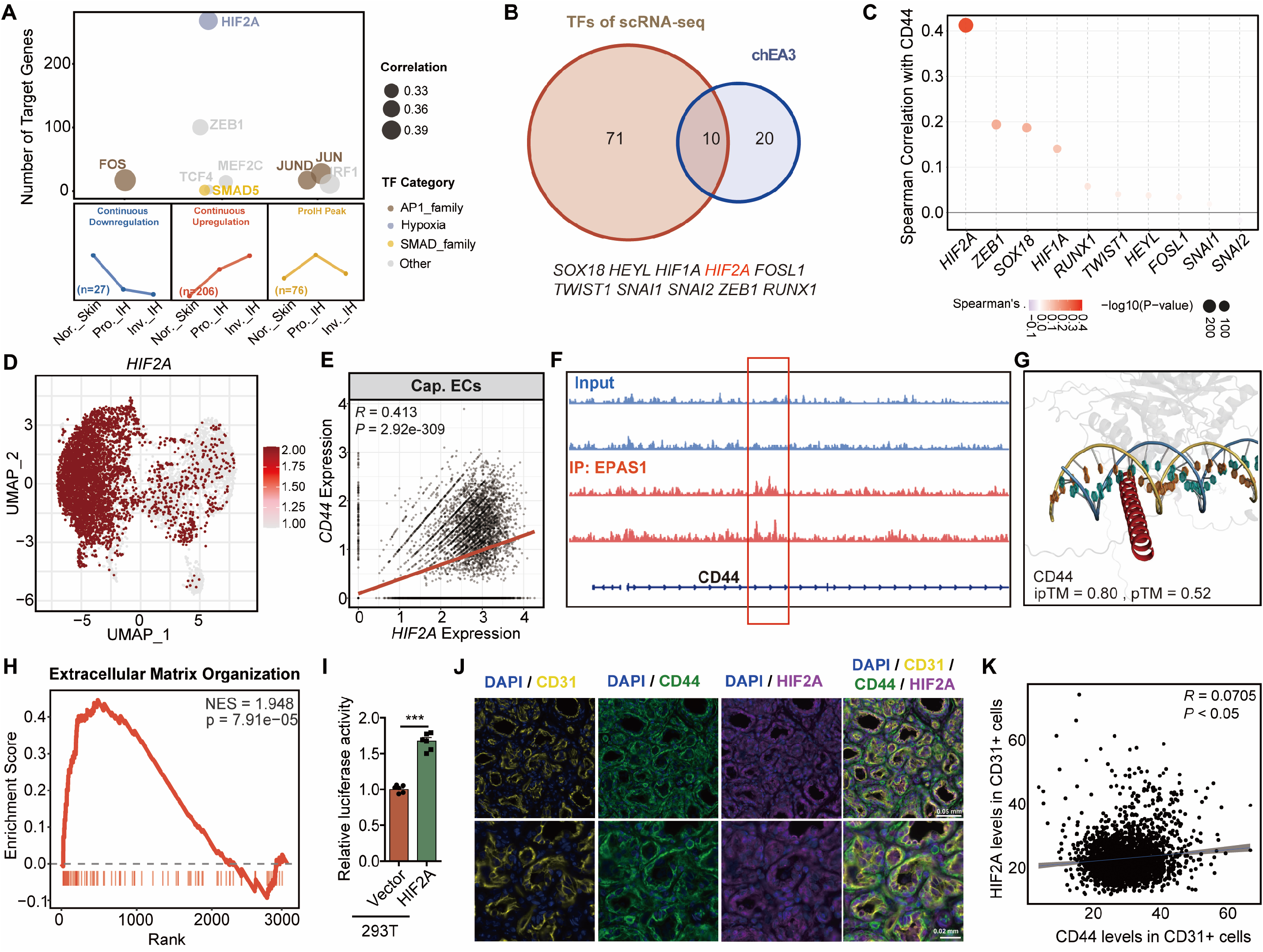
HIF2A activates the transcription of CD44 in Cap.ECs. (A) Temporal expression of TFs in Nor.Skin, Pro.IH, and Inv.IH. (B) Overlapping expression of ten TFs in the CD44 signaling pathway (including HIF2A) and in proliferative Cap.ECs. (C) Correlation of the expression of ten TFs with CD44 expression in Cap.ECs. (D) UMAP plot showing the differential expression of HIF2A in different subpopulations of vascular endothelial cells. (E) Correlation in the expression of HIF2A with CD44 in Cap.ECs. (F) Public ChIP-seq datasets reveal a distinct HIF2A binding peak at the CD44 promoter region. (G) Structural modeling, demonstrating high binding affinity of the HIF2A protein and the CD44 DNA motif. (H) In silico (virtual) knockout of HIF2A, showing severe disruptions in the organization of the ECM signaling. (I) Dual-luciferase reporter assays showing the transcriptional activity of the CD44 promoter in 293T with HIF2A overexpression. (J) Representative multiplex immunofluorescence images of tissue sections, showing co-localization of HIF2A (green) and CD44 (red). DAPI (blue) was used for nuclear counterstaining. Scale bar: 50 μm. (K) Results of linear correlation between fluorescence quantification of CD44 and HIF2A.

We further examined this topic by performing *in vitro* studies with a vector that overexpresses *HIF2A* in 293T. The results from dual-luciferase reporter assays confirmed that *HIF2A* overexpression enhanced the transcriptional activity of the CD44 promoter (Fig. 4I). Multicolor *in situ* staining of clinical tissues also showed a strong cellular co-localization of HIF2A and CD44 during the proliferation phase (Fig. 4J, K). Together, these findings indicated that HIF2A is a key TF that increases the expression of CD44 in Cap.ECs. These results provide a link of microenvironmental hypoxia to robust angiogenic responses during the proliferation phase of IH.

### *SERPINE2^+^* pericytes sustain the angiogenesis of Cap.ECs *via* ANGPT signaling

The identification of pericytes as the most frequent neighbors of Cap.ECs led us to examine the mechanism by which pericytes regulate the endothelial cells. We then used scRNA-seq profiling to investigate pericyte-mediated regulation by classification of pericytes into six distinct subpopulations (Fig. 5A, B). Angiogenic and *SERPINE2^+^* pericytes were more abundant during the proliferation phase (Fig. 5B, Supplementary Table 7), and were characterized by increased signaling of the PI3K-AKT, VEGF, and hypoxia pathways (Fig. 5C). In contrast, inflammatory pathways were more active during the involution phase (Fig. 5B, C, Supplementary Table 8). Pseudotime analysis revealed that, during the proliferation phase, cells clustered at the origin and early branches (Angiogenic and *SERPINE2^+^* pericytes); during the involution-phase, cells shifted to the middle and late branches and were mature and inflammatory pericytes (Supplementary Fig. 9A, B). Angiogenic signaling pathways and key genes were initially highly active, but decreased over time, which is consistent with the temporal trajectory of proliferative-phase cells (Supplementary Fig. 9B-D). These findings highlight a critical shift in the phenotype of pericytes during the transition from proliferation to involution, which is paralleled by a shift in angiogenic signaling activity.

**Figure 5.**
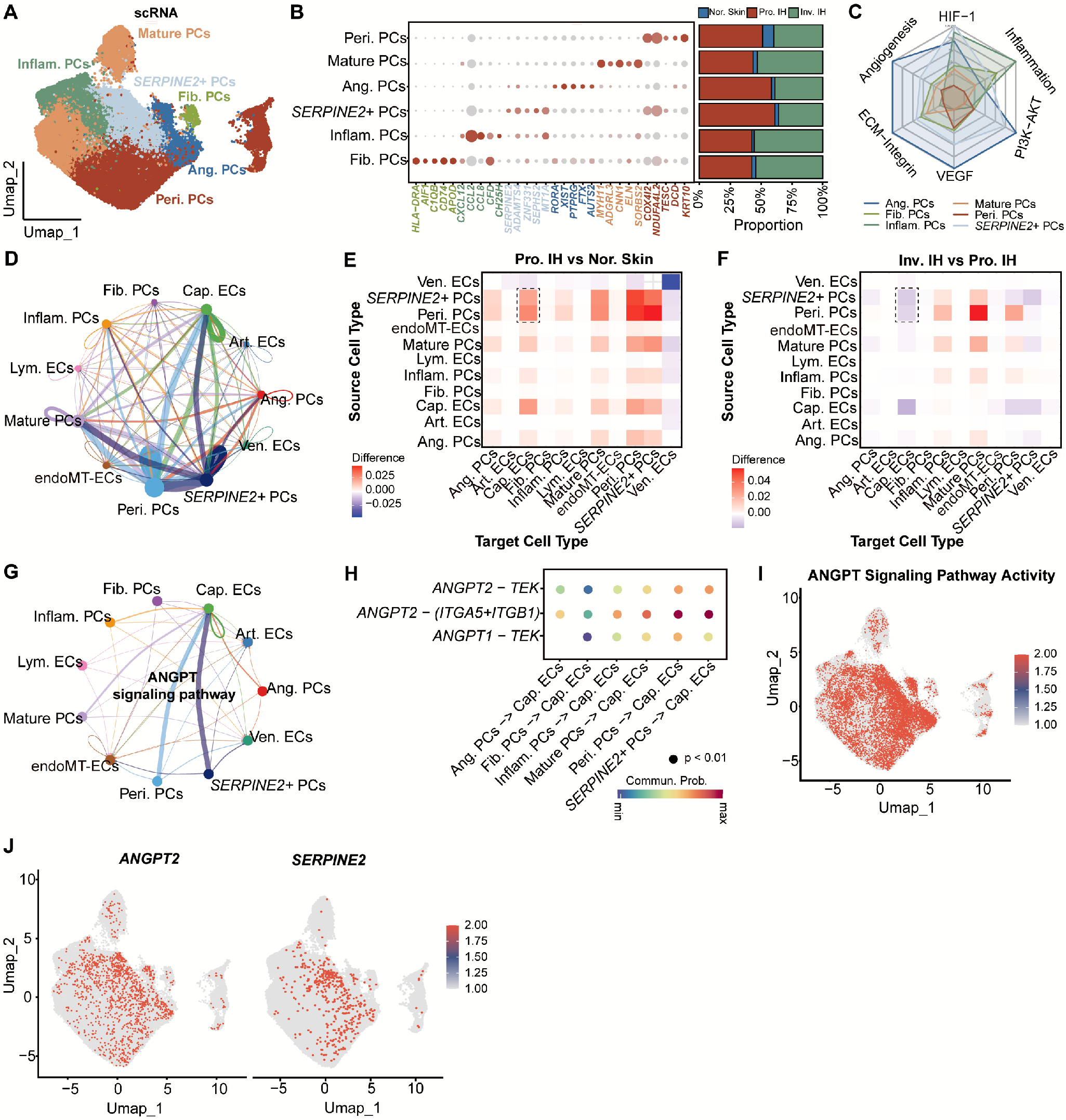
*SERPINE2+* pericytes sustain the angiogenesis of Cap.ECs via ANGPT signaling. (A) UMAP plot showing the re-clustering of 67,546 pericytes into six distinct subpopulations. (B) Expression of canonical markers for each pericyte subgroup (left), and proportions of pericyte subpopulations in Nor.Skin, Pro.IH, and Inv.IH (right). (C) Pathway enrichment scores for each pericyte subpopulation. (D) Circular plot depicting the communication strength between pericyte subpopulations and endothelial cell subpopulations in the Pro.IH group. (E) Relative strength of communication of each subpopulation of pericytes with Cap.ECs in Pro.IH vs. Nor.Skin (scRNA-seq data). (F) Relative strength of communication of each subpopulation of pericytes with Cap.ECs in Inv.IH vs. Pro.IH (scRNA-seq data). (G) Ligand-receptor pair analysis of ANGPT signaling in the communication of each pericyte subpopulation with Cap.ECs. (H) Expression of significant ligands in the different pericyte subpopulations (scRNA-seq data). (I) Feature plots depicting the distribution and intensity of ANGPT signaling activity among various pericyte subpopulations. (J) Feature plots depicting the distribution and intensity of ANGPT2 and SERPINE2 among various pericyte subpopulations.

Interactome analysis further revealed that in addition to classical pericytes (Peri.PCs), *SERPINE2^+^* pericytes were the primary drivers of Cap.EC-mediated angiogenesis during Pro.IH (Fig. 5D-F), mainly by increased signaling *via* the ANGPT pathways (Fig. 5G, H). Notably, *SERPINE2+* pericytes had upregulation of genes encoding messengers (such as *ANGPT2*) in this pathway (Fig. 4I, J), suggesting that these cells function in the sustained activation of Cap.ECs angiogenesis. Additionally, *ANGPT2* shares similar cellular localization with *SERPINE2* (Fig. 4J), and their expression is positively correlated (Fig. 4K), further implying specific *ANGPT2* expression in *SERPINE2⁺* pericytes. Together, these data indicate the presence of interactions of pericytes and Cap.ECs in IH tissues, in which *SERPINE2⁺*pericytes stabilize and promote Cap.EC-mediated vascular expansion via ANGPT signaling during the proliferation phase. Thus, targeting *SERPINE2* or ANGPT signaling may offer a novel therapeutic avenue for proliferating IH, potentially complementing existing anti-angiogenic strategies.

### MSCs enriched niche provides a progenitor pool for Cap.ECs during the proliferation phase

Because Cap.ECs have a key role in driving angiogenesis, we examined the cellular origins of these cells. Analysis of the pseudotime trajectory revealed a distinct developmental continuum, in which immature Cap.ECs in the proliferation phase differentiated from highly multipotent MSCs (Fig. 6A). In contrast, this transition was not present in normal skin and involuting IH, and both lineages remained in mature and stable states in these other tissues (Fig. 6A). Within the highly angiogenic N8 (MSCs), spatial distance mapping confirmed that MSCs were adjacent to Cap.ECs (Fig. 6B-D). Moreover, there was evidence of crosstalk between these two populations (Fig. 6E), consistent with spatial and temporal interactions during early differentiation.

**Figure 6.**
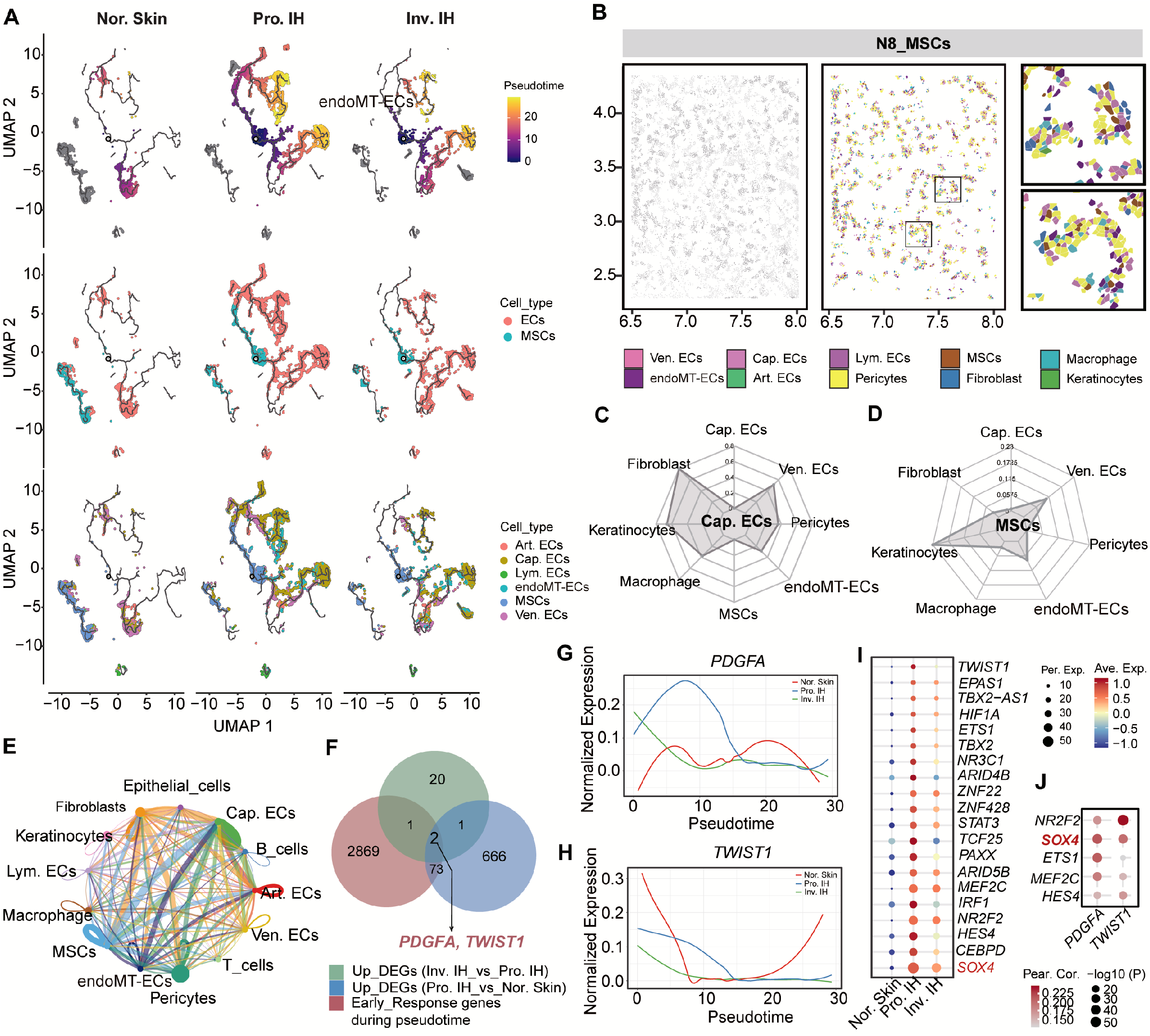
MSCs enriched niche provides a progenitor pool for Cap.ECs during the proliferation phase. (A) Pseudotime trajectory analysis of the distinct developmental continuum, in which highly multipotent MSCs differentiate into immature Cap.ECs in Pro.IH. (B) Spatial pattern of N8 (MSCs) in serial tissue sections from Pro.IH. (C) Mean distance (axis length) of different types of cells to Cap.ECs. (D) Mean distance (axis length) of different types of cells to MSCs. (E) Cell chart analysis of the single-cell interactome, demonstrating robust crosstalk between MSCs and Cap.ECs. Line thickness represents strength of crosstalk. (F) Intersection of genes enriched in early pseudotime states with those highly expressed in Pro.IH MSCs, indicating PDGFA and TWIST1 as critical lineage-commitment regulators. (G, H) Expression of PDGFA and TWIST1 during the initial stages of the pseudotime trajectory, showing a gradual decline as differentiation progresses toward mature endothelial cells. (I) Thirty TFs with the highest expression in MSCs of Pro.IH. (J) Correlation of the levels of PDGFA and TWIST1 with the top 3 TFs in MSCs of Pro.IH.

To identify the molecular drivers of the transition from MSCs to Cap.ECs, we compared genes enriched in early pseudotime states with those enriched in the proliferation phase. This unbiased approach indicated that *PDGFA* and *TWIST1* were key regulators of this process (Fig. 6F). The expression of both genes peaked at an early pseudotime, and expression gradually declined as differentiation progressed toward mature HemECs (Fig. 6G, H). Differential expression analysis of the MSC population also showed that SOX4 regulated the PDGFA and TWIST1 cascade (Fig. 6I, J). Together, these findings indicate that hypoxia-driven MSCs are the primary progenitors of Cap.ECs, and that they promote pathological angiogenesis during the early proliferation phase of IH. In addition, the hypoxic microenvironment drives the differentiation of MSCs into HemECs during the early proliferation phase of IH, and SOX4-regulated expression of PDGFA and TWIST1 regulate lineage commitment.

## DISCUSSION

IH is characterized by an excessive but self-limiting angiogenesis [9]. Although previous studies have identified the composition of cells in IH, no previous studies have characterized the spatiotemporal organization of the microenvironment and the intercellular interactions that underlie cell proliferation. Our integration of scRNA-seq, high-resolution spatial transcriptomics, and functional validation enabled a detailed characterization of IH tissues during the proliferation and involution phases. Our results indicated that hypoxia triggered the transition of MSCs into HemECs and also activated these endothelial cells *via* the HIF2A/CD44 signaling axis. We also identified reprogrammed pericytes as the central regulators that modulate the function of Cap.ECs. These dynamic interactions and regulators, such as SERPINE2 (a serine protease inhibitor), lead pericytes and Cap.ECs to drive the pronounced vascular malformation and tissue remodeling that is characteristic of the proliferation phase of IH.

Our spatially resolved single-cell atlas showed that HemECs adopted a highly complex, quasi-neoplastic phenotype during the proliferation phase. Beyond the well-documented dependence of HemECs on VEGF [10], HemECs exhibit robust hyperactivation of processes classically linked to cancer progression during the proliferation phase, including the PI3K-AKT pathway [11, 12], GTPase activity [13], and ECM remodeling [14]. Notably, the upregulation of embryogenesis-related genes (DLK1) and markers of tumor stemness (ROR1) suggests that Cap.ECs recapitulate developmental angiogenesis [15–17]. This spatial and transcriptomic evidence identified IH proliferation as an aggressive stage of initiation, in which the tumor-like metabolic and growth of HemECs increase vascular expansion.

Previous studies have had disparate conclusions regarding the cellular origin of the massive expansion of endothelial cells during the pathogenesis of IH. Early studies proposed a placental origin or involvement of circulating endothelial progenitor cells (EPCs) [18], and later studies emphasized multipotent hemangioma stem cells (HemSCs) that are capable of forming vascular structures [19]. However, these models were largely based on *in vitro* findings, which prevented consideration of the spatial distribution of cells in the TME. Our study provides direct *in situ* evidence that local MSCs are the progenitors of Cap.ECs, and characterization of spatial transcriptomics identified the trans-differentiation from multipotent progenitors (MSCs) to ECs to drive the pathogenesis of IH [20, 21]. The presence of this transition during the proliferation phase, but not in normal skin or during the involution phase, indicates that this is a highly localized and disease-specific pathological event. Although hypoxia is accepted as a key driver of cell proliferation, the molecular mechanism by which it determines the fate of stem cells remains unclear [24, 25]. Our results indicated that a previously unrecognized transcriptional cascade, in which the hypoxia-induced factor SOX4 [26, 27] directly regulates the expression of PDGFA and TWIST1, triggers the lineage commitment of MSCs. Although TWIST1 is regarded as a master regulator of the epithelial-mesenchymal transition in tumor metastasis [28, 29], we demonstrated it also drives the mesenchymal-to-endothelial transition (MEndoT) during pathological angiogenesis. Our characterization of the SOX4-PDGFA/TWIST1 axis demonstrated that the hypoxic microenvironment reprograms the local MSCs and actively supplies a pool of progenitor cells required for increased vascular growth.

Pericytes are typically quiescent mural cells that provide structural support to mature endothelial networks [30–32]. However, our high-resolution spatial and single-cell analyses indicated they have an active role in the pathogenesis of IH. Rather than functioning as passive stabilizers, pericytes in the proliferation phase of IH functioned as highly dynamic and active drivers of vascular expansion. We identified significant communication between proliferative pericytes and Cap.ECs during the proliferation phase. This is likely to be a critical step in promoting angiogenesis and vascular expansion, hallmarks of the early phase of IH [33]. Beyond intercellular communication, our results identified a profound intracellular functional reprogramming within the proliferative pericytes that closely mirrored the stromal adaptations in a malignant TME [34, 35]. Although previous research has implicated SERPINE2 in remodeling of the ECM and tumor progression in various cancers [36, 37], our identification of its role in regulating pericyte plasticity and secretion of pro-angiogenic ligands (ANGPT2, PGF) in IH is a novel finding.

A central finding of our study is the identification of a non-canonical COL4A1/2–CD44 signaling axis as the primary mediator of crosstalk between Peri.ECs and Cap.ECs during proliferative IH. Although CD44 is known as a hyaluronan receptor [38], its role in transducing collagen IV signals between subsets of endothelial cells in IH is a novel finding. Our spatial distance analysis indicated a close physical proximity of these cells, and this likely enabled a robust paracrine feedback loop, in which matrix-producing Peri.ECs deposit type IV collagen that directly engages CD44 on neighboring Cap.ECs. The functional significance of this signaling extends beyond mere structural support, because it acts as a critical mechanotransducer that integrates ECM cues with intracellular angiogenic programs [39, 40]. This functional evidence aligns with our another finding that HIF2A regulates the transcription of CD44 during hypoxic stress, and underscores HIF2A as the predominant hypoxia-inducible factor in IH [41]. Unlike HIF1A (which primarily governs acute hypoxic responses), HIF2A is closely linked to chronic hypoxia and plays a central role in sustaining vascular proliferation and endothelial cell survival in IH [42, 43]. Previous research showed that HIF2A regulates the expression of VEGF and other angiogenic factors, thereby contributing to the increased vascular growth that is characteristic of IH [44]. Targeting HIF2A and its downstream signaling pathways is a potential therapeutic approach that may slow the progression or promote the regression of IH.

Although this study provided a comprehensive single-cell and spatial transcriptomic atlas of IH that established a new framework for understanding the pathogenesis of this tumor, several limitations warrant consideration. First, although our multi-omic approach provided novel insights, the findings were mostly descriptive. Identification of causal relationships within the signaling networks identified here requires definitive validation using gain- and loss-of-function experiments with xenograft or other *in vivo* models. Second, although our spatial transcriptomics provided critical topographical context, higher-resolution technologies are needed to validate the predicted ligand-receptor interactions at single-cell resolution. Moreover, our study had certain technical limitations related to tissue dissociation, patient heterogeneity, and the absence of multi-omic integration (epigenomics or proteomics). Finally, the therapeutic efficacy of targeting the molecular nodes identified here has yet to be evaluated in preclinical models. Ultimately, there is a need for clinical validation in independent, large cohorts with longitudinal follow-up. Addressing these limitations in mechanistic studies that examine the pathogenesis of IH and in clinical studies that perform longitudinal follow-up will be essential for translating the novel findings of the present study into novel diagnostic and therapeutic strategies for patients with IH.

In conclusion, our integrated single-cell and spatial transcriptomic atlas showed that a highly coordinated and spatiotemporally dynamic environment drives the pathogenesis of IH (Fig. 7). We showed that vascular growth during the proliferation phase of IH is driven by the specific niche centered around Cap.ECs. Hypoxia-primed MSCs continuously seed this pathological niche, and the dual pro-angiogenic and immunosuppressive Cap.EC phenotype sustains the niche. We also identified strong spatial interactions during the progression of IH. In particular, Cap.ECs were structurally and molecularly reinforced by transitional endoMT-derived endothelial cells through a matrix-driven COL4A1/2-CD44 paracrine loop and by *SERPINE2^+^* pericytes due to extensive bidirectional signaling. Our analysis of these complex multicellular interactions reframes the developmental origins and spatial organization of IH, and also suggests that microenvironmental crosstalk may be a vulnerability that can potentially be exploited by novel therapeutics. Disrupting these matrix-mediated interactions, particularly blockage of CD44 signaling in combination with standard propranolol treatment, has potential as a potent, mechanistically grounded approach to disrupt the pathological vascular network of IH and achieve durable tumor regression.

## MATERIALS AND METHODS

### Collection and processing of clinical samples

This study was approved by the Institutional Ethics Review Committee of Jinan Children’s Hospital (Approval No.: SDFE-2026038) and was conducted in strict accordance with the Declaration of Helsinki. Written informed consent was obtained from the legal guardians of all participants. A total of 30 pediatric tissue samples were collected from patients undergoing surgical resection and were divided into three groups: proliferative-phase infantile hemangioma (Pro.IH, age 0–6 months), involuting-phase infantile hemangioma (Inv.IH, age 1–2 years), and age-matched normal skin (Nor.Skin, age 1–2). Two pathologists independently confirmed the diagnosis and phase of disease in each patient. The baseline clinical data, including age, gender, and lesion characteristics, were anonymized and prospectively recorded for blinded downstream analysis. Immediately after surgical resection, tissues were allocated and processed for specific downstream analyses. For scRNA-seq, fresh tissue blocks (∼0.5 cm^3^) were submerged in pre-cooled MACS™ Tissue Storage Solution (Miltenyi Biotec) at 4 °C and transported to the laboratory for single-cell dissociation within 2 h; if processing was delayed, samples were cryopreserved in a freezing medium (90% fetal bovine serum + 10% DMSO) and stored in liquid nitrogen. For spatial transcriptomics, fresh tissues were formalin-fixed and paraffin-embedded (FFPE). Then, guided by H&E staining, representative regions were selected and assembled into a 1 × 1 cm tissue microarray (TMA). For multiplex immunofluorescence (mIF), serial 5 μm sections were cut from the FFPE blocks, mounted onto SuperFrost Plus slides (Thermo Fisher), baked at 60 °C for 2 h, and then stored at room temperature until staining.

### Single-cell RNA sequencing and data processing

Fresh tissues (6 Pro.IH, 4 Inv.IH, and 5 Nor.Skin) were minced and enzymatically digested with collagenase IV (1 mg/mL, Worthington) and DNase I (0.1 mg/mL, Sigma) at 37 °C for 30 min. After confirming viability (>85% by trypan blue exclusion), single-cell libraries were constructed using the Chromium Single Cell 3′ Reagent Kit (ver. 3, 10x Genomics). The libraries were sequenced on an Illumina NovaSeq 6000 platform to a target depth of 100,000 reads per cell.

Raw sequencing reads were processed with the Cell Ranger pipeline (ver. 6.0, 10x Genomics) and aligned to a human reference genome (GRCh38). To ensure data quality, cells with fewer than 200 detected genes or more than 20% mitochondrial transcripts were excluded. Doublets were identified and removed using DoubletFinder (ver. 2.0); the expected doublet rate was ∼5.2%. After rigorous quality control, data from 172,683 high-quality cells were retained for downstream analysis.

Downstream analyses were conducted using Seurat (ver. 5.0.0). Gene expression matrices were log-normalized, and the 3,000 most highly variable genes (HVGs) were selected for principal component analysis (PCA). To correct for batch effects among the 15 samples while preserving biological variation, the Harmony algorithm (ver. 0.1.0) was applied to the PCA embeddings.

Graph-based clustering was performed on the first 30 Harmony-corrected dimensions using the Louvain algorithm (resolution = 0.8). Cell types were annotated according to the following canonical markers: endothelial cells (CD34, PECAM1, KDR), pericytes (RGS5, ACTA2, PDGFRB), fibroblasts (LUM, DCN), T cells (CD3D, CD3E), macrophages (CD163, CSF1R, CD14), keratinocytes (KRT5, KRT10, KRT1), epithelial cells (EPCAM, KRT8, KRT18), B cells (CD79A, IGKC), and MSCs (MKI67, UBE2C, TOP2A). Finally, cluster-specific marker genes were identified using the Wilcoxon rank-sum test. Standard significance thresholds were used for comparisons (adjusted *p* < 0.05 and |log_2_ fold change| > 0.25).

### Spatial transcriptomics (CosMx® SMI) and niche identification

Tissue microarrays (TMAs) from 399 fields of view (FOVs) were constructed from 15 FFPE samples (4 Pro.IH, 6 Inv.IH, and 5 Nor.Skin). Then, 5-μm sections were mounted onto customized slides. Spatial transcriptomics was performed using the CosMx® Spatial Molecular Imager (SMI, NanoString Technologies) with a custom 1,000-gene panel, strictly following the manufacturer’s protocol.

Raw imaging data were processed using the CosMx® Data Analysis Suite (ver. 1.5). Single cells were segmented using DAPI (nucleus) and antibodies against pan-cytokeratin and CD45 (membrane). Following segmentation and quality control, 634,558 individual cells were identified and segregated into major cell lineages based on the spatial pattern of gene expression. Spatial transcriptomics data were processed using Seurat and Giotto (ver. 1.0). Raw spatial expression matrices were log-normalized, and low-quality spots (<100 detected genes or >20% mitochondrial content) were excluded.

To resolve cellular heterogeneity at near-single-cell resolution, spatial spots were deconvoluted using SPOTlight with non-negative matrix factorization. Cell type-specific signatures derived from the scRNA-seq data were used as references to map the five distinct subtypes of endothelial cells. Then, spatial probability scores were generated for each spot to enable precise quantification of the distribution of different subtypes of endothelial cells and their abundances in the different phases of IH. To systematically define recurrent tissue microenvironments, unsupervised *k*-means clustering was applied to the integrated cellular composition matrix of each spot (combining deconvoluted proportions of endothelial cells with the abundances of other major lineages). The optimal number of clusters (k = 10) was determined using the elbow method and silhouette score analysis and indicated the tissue architecture had ten distinct spatial niches.

### Differential gene expression and functional enrichment analysis

Differentially expressed genes (DEGs) were identified using the Seurat FindMarkers function (Wilcoxon rank-sum test), with significance thresholds set at standard values (adjusted *p* < 0.05 and |log_2_ fold change| > 1). To characterize gene expression in endothelial cells in Nor.Skin, Pro.IH, and Inv.IH, genes were grouped into three distinct clusters based on temporal expression. Functional enrichment using Gene Ontology (GO) biological processes and Kyoto Encyclopedia of Genes and Genomes (KEGG) pathways was performed using the R package *clusterProfiler*, and an adjusted *p*-value below 0.05 was considered significant.

### Pseudotime and trajectory analysis

To objectively establish the developmental hierarchy of endothelial cells, cellular stemness and differentiation potential were first evaluated using CytoTRACE (standard stemness scoring). Pseudotime cellular trajectories were then inferred with the Monocle3 package (ver. 1.0). Because they had the highest stemness scores and had proliferative- and progenitor-like characteristics, Cap.ECs were considered to be the root state of the differentiation trajectory.

### Analysis of intercellular communications

Intercellular communication networks were analyzed using CellChat (ver. 1.1.3). Ligand-receptor interactions were inferred by determining the expression of ligands and receptor genes in defined cell populations. For each ligand-receptor pair, communication strength and probability were calculated for normal skin, the proliferation phase, and the involution phase. Significant interactions were identified using permutation tests with 1,000 iterations. The key signaling pathways — COLLAGEN, VEGF, PECAM1, and ANGPT-TEK — were visualized using circle plots and heatmaps.

### Multiplex immunofluorescence staining

FFPE tissue sections (4 μm) were deparaffinized, rehydrated, and subjected to antigen retrieval in a citrate buffer (pH 6.0) at 95 °C for 20 min. After blocking with 10% normal donkey serum, the sections were incubated overnight at 4 °C with the following antibodies: anti-CD31 (Cat No. 66065-2-Ig, Proteintech), anti-RGS5 (Cat No. 11590-1-AP, Proteintech), anti-CD44 (Cat No. 60224-1-AP, Proteintech), anti-COL4A1 (Cat No. 30850-1-AP, Proteintech), anti-NOX4 (Cat No. 67681-1-AP, Proteintech), anti-ROR1 (Cat No. 66923-1-AP, Proteintech), anti-PDGFRB (Cat No. 13449-1-AP, Proteintech), and anti-RGCC (Cat No. 19556-1-AP, Proteintech). Detection was performed using secondary antibodies conjugated to Opal fluorophores (Akoya Biosciences). Slides were counterstained with DAPI and viewed using the Vectra Polaris multispectral imaging system (Akoya Biosciences). Cell densities and spatial proximity were quantified using inForm software (ver. 2.4, Akoya) and HALO (Indica Labs).

### Isolation and culture of endothelial cells from primary hemangioma

Fresh tissues were minced and digested with 2 mg/mL collagenase II (Worthington) at 37 °C for 1 h. The cell suspension was passed through a 40 μm strainer, and CD31+ endothelial cells were isolated by magnetic-activated cell sorting (MACS) using CD31 microbeads (Miltenyi Biotec). The isolated HemECs were cultured in ECM medium (ScienCell) containing 5% fetal bovine serum on fibronectin-coated plates (10 μg/mL, Sigma). Human umbilical vein endothelial cells (HUVECs) were obtained from ScienCell and were maintained in ECM medium. All cells were maintained at 37 °C in a humidified incubator with 5% CO_2_.

### *In vitro* angiogenesis and co-culture assays

For tube formation assays, 48-well plates were coated with 100 μL of Matrigel (Corning) per well and allowed to polymerize at 37 °C for 30 min. HUVECs or HemECs were then seeded at 5 × 10⁴ cells per well and incubated for 6 to 8 h. Tube formation was quantified by measuring total tube length and branch points using ImageJ software. In co-culture experiments, HemECs were seeded in transwell inserts (0.4 μm pores, Corning) and co-cultured with HUVECs in the bottom chamber for 24 to 48 h prior to assessment of angiogenesis. For CD44 inhibition experiments, cells were pretreated with an anti-CD44 neutralizing antibody (RO-5429083, 0.5 μg/mL) or an isotype control for 1 h before seeding.

### Plasmids and construction of stable cell lines

To construct cells that had stable overexpression of *HIF2A*, the coding region of *HIF2A* was subcloned into the pLV3-CMV-MCS-3xFLAG-CopGFP-Puro lentiviral overexpression vector. The resulting plasmids were co-transfected into 293T cells with psPAX2 and pMD2.G at a ratio of 4:3:1 to produce lentiviral particles. HemECs were then infected with the lentiviral particles. After 48 h, the cells were treated with 2 μg/mL puromycin and selected for at least four days. Finally, HIF2A expression in the cells was determined by western blotting using an anti-HIF2A monoclonal antibody (#66731-1-Ig, Proteintech, China).

### Western blotting

Cells were lysed on ice for 30 min using RIPA lysis buffer (#sc-24948, Santa Cruz) that contained 1 mM PMSF (#P7626, Sigma) and phosphatase inhibitors. The lysates were centrifuged at 13,000 *g* for 15 min at 4 °C to isolate proteins. Protein concentration was measured with a bicinchoninic acid (BCA) assay kit (#P0009, Beyotime, China). Thirty micrograms of protein from each sample was loaded into each well of a 10% SDS-PAGE gel, and proteins were separated by electrophoresis at 140 V for 60 min. The proteins were transferred onto a 0.45 μm PVDF membrane, blocked with 5% non-fat dry milk for 1 h at room temperature, and incubated overnight at 4 °C with a primary monoclonal antibody against CD44 (#60224-1-Ig, Proteintech, China). The membrane was then incubated with an HRP-linked secondary antibody (#SA00001-1, Proteintech, China) for 1 h at room temperature, and developed using the Immobilon Forte Western HRP substrate (#E411, Vazyme, China). Images were acquired with an ultrasensitive multifunctional fluorescence chemiluminescence imaging system (Chemiscope 6200Touch, Shanghai Qinxiang Scientific Instrument Co., LTD, China).

### Detection of double luciferase reporter gene activities

The promoter sequence (Chr.11:35136882-35138981) of the *CD44* gene was subcloned into the pGL3-basic vector, and was linked to the luciferase sequence. The pGL3-Basic-pCD44-luciferase plasmid (or the empty vector) was transfected into HemECs that overexpressed *HIF2A* (or control vector), along with the TK plasmid, and cells were then cultured for 24 h. These cells were treated with the lysis buffer from the dual-luciferase reporter assay kit (RG027, Beyotime, China). Following complete lysis, the lysates were transferred to a 1.5 mL centrifuge tube and centrifuged (10,000–15,000 *g* for 3–5 min). The supernatant was collected and the relative light units (RLU) of firefly and *Renilla* luciferase were measured using a Spark Multimode Microplate Reader (TECAN SPARK, Switzerland).

### Animal studies

All animal experiments were approved by the Institutional Ethics Review Committee of Ji’nan Children’s Hospital (Approval No.: SDFE-2026038). For the hemangioma mouse model, HemECs were mixed with HUVECs at a 1:1 ratio (1 × 10^6^ cells total) in Matrigel, and were then injected subcutaneously into the flanks of 6-week-old female BALB/c nude mice (6 per group). Tumor growth was monitored every 3 days by caliper measurements. At 3 days after injection, mice received vehicle control, an anti-CD44 neutralizing antibody (RO-5429083, 1 mg/kg, intraperitoneal injection daily), propranolol (40 mg/kg, daily oral gavage), or combination therapy. On 7 days after injection, the mice were euthanized and tumors were harvested for measurement of tumor volume ([length × width^2^]/2) and histological analysis.

### In silico methods

Single-cell data were loaded from a pre-annotated Seurat object, and endothelial cells (n = 14,464) were extracted. Raw counts were filtered to retain genes expressed in at least 5% of cells, and mitochondrial (MT-) and ribosomal (RPS/RPL) genes were then removed. Genes were ranked by the coefficient of variation, and the 3,000 genes with the greatest variation were selected, along with *EPAS1*. *EPAS1* knockout was simulated using scTenifoldKnk with the following parameters: nc_nNet = 7, nc_nCells = 400, nc_nComp = 3, nc_q = 0.92, td_K = 3, ma_nDim = 2, and nCores = 16. Pathway analyses were performed with enrichR (GO BP, KEGG, Hallmark, WikiPathways), and GSEA was conducted using fgsea, with genes ranked by descending *z*-score. Figures were generated using the *ggplot2* package in R.

### CHIP-seq analysis

The 786-O ChIP-seq data were obtained from the NCBI GEO/SRA database (BioProject accession number: PRJNA494833). Raw FASTQ sequencing reads were initially assessed for quality using FastQC. Adapter trimming and quality filtering were then conducted with fastp using the following parameters: -q 20, -u 20, and -l 20. Clean reads were aligned to a human reference genome GRCh38 (hg38) using Bowtie2, and the resulting alignments were sorted and converted to BAM format using samtools. PCR duplicates were removed using sambamba markdup. Peak calling was performed with MACS2 (*p*-value cutoff: 0.05) and the hs genome size parameter. Each immunoprecipitation sample was matched against its corresponding Input control. Peaks were annotated to genomic features relative to the transcription start site (TSS ± 3 kb) using the R package *ChIPseeker*. The, *de novo* and known motif enrichment analyses were performed on merged peak regions (± 100 bp) using findMotifsGenome.pl in the HOMER suite. Based on motif analysis, DNA sequences surrounding the TSSs were extracted. The AlphaFold Server (https://alphafoldserver.com) was subsequently used to predict the structural complex and binding interaction between TFs and target DNA sequences. For visualization, deepTools (bamCoverage, computeMatrix, and plotHeatmap) was employed to generate BigWig signal tracks and produce heatmaps of ChIP-seq signal density around the TSSs.

### Statistical analysis

All statistical analyses and data visualizations were performed using R (ver. 4.0) and GraphPad Prism (ver. 9.0). Quantitative data are expressed as mean ± standard error of the mean (SEM) unless otherwise stated. Comparisons of two independent groups were performed using a two-tailed Student’s *t*-test. Comparisons of three or more groups were performed using a one-way analysis of variance followed by Tukey’s *post-hoc* test. Linear correlations were determined by calculation of Pearson’s correlation coefficient. For bioinformatics analyses, differential gene expression in single-cell and spatial transcriptomic data were assessed using the Wilcoxon rank-sum test, with Bonferroni correction for multiple testing. A *p*-value below 0.05 was considered significant and the figures indicate the level of significance as * (*p* < 0.05), ** (*p* < 0.01), and *** (*p* < 0.001).

## Supporting Information

Supporting Information is available from the Wiley Online Library or from the author.

## Data Availability

All data are provided in the manuscript and in the supplementary materials. The raw data generated in this study, including including single-cell RNA sequencing data, CosMx spatial molecular imaging data, bulk RNA sequencing data of IH tissue, and bulk RNA sequencing data of HemECs, have been deposited in the NGDC database under accession number HRA018919 (BioProject ID: PRJCA065790).

## Acknowledgements

This work was supported by the Shandong Provincial Natural Science Foundation (ZR2022MH236 to L.G., and ZR2023QH305 to X.G.), Shandong Province Medical and Health Science and Technology Development Project (202506020863 to L.G.), Research Projects Approved by the National Health Commission Capacity Building and Continuing Education Center in 2023(GWJJ2023100303 to L.G.), The Special Fund Project for Industrial Leading Talents of the Haiyou Plan (Ji Rencai Ban Fa (2024) 5 to L.G.), Jinan Health High-Caliber Talent Project (202312 to L.G.).

## Conflict of Interest

The authors declare no conflict of interest.

## Author Contributions

X.G. conceived the concept, carried out experiments, prepared figures and tables, designed and wrote the manuscript, and was responsible for the corresponding works. Z.L., and L.Y. were responsible for part of the bioinformatics analysis, and participated in guiding the project and critically revised the manuscript. Z.C., and D.X. were responsible for sample collection and animal experiments, and critically revised the manuscript. L.W, P.Y., B.Z., Y.N., L.Y., and J.H. participated in partial experiments and critically revised the manuscript. L.W., was responsible for the supervision and resection of clinical samples, as well as data organization. H. Z., and K. Z. critically revised the manuscript. D.S. was responsible for the supervision and resection of clinical samples and supervised results. Z.L. coordinate the entire project, supervised results, designed and critically revised the manuscript, and was responsible for the corresponding works. L.G. conceived the concept, coordinate the entire project, supervised results, designed and critically revised the manuscript, and was responsible for the corresponding works. All authors approved the final manuscript.

## Notes

### Competing Interest Statement

The authors have declared no competing interest.

